# Blood RNA Profiles are Diagnostic for Severity in Human Acute Spinal Cord Injury

**DOI:** 10.1101/2020.04.15.037325

**Authors:** Nikos Kyritsis, Abel Torres Espin, Patrick G. Schupp, J. Russell Huie, Austin Chou, Xuan Duong-Fernandez, Leigh H. Thomas, Rachel E. Tsolinas, Debra D. Hemmerle, Lisa U. Pascual, Vineeta Singh, Jonathan Z. Pan, Jason F. Talbott, William D. Whetstone, John F. Burke, Anthony M. DiGiorgio, Philip R. Weinstein, Geoffrey T. Manley, Sanjay S. Dhall, Adam R. Ferguson, Michael C. Oldham, Jacqueline C. Bresnahan, Michael S. Beattie

## Abstract

Biomarkers of spinal cord injury (SCI) could help determine the severity of the injury and facilitate early critical care decision making. We analyzed global gene expression in peripheral white blood cells during the acute injury phase and identified 197 genes whose expression changed after SCI compared to healthy and trauma controls and in direct relation to SCI severity. Unsupervised co-expression network analysis identified several gene modules that predicted injury severity (AIS grades) with an overall accuracy of 72.7% and included signatures of immune cell subtypes. Our findings indicate that global transcriptomic changes in peripheral blood cells have diagnostic and potentially prognostic value for SCI severity.

## INTRODUCTION

Precision medicine (often interchangeably called personalized medicine) promises to optimize individualized treatment options based on demographic and genetic characteristics as well as the specific biological features of the presenting disorder. Cancer therapy has already benefited greatly from this approach,^1,2^ where blood- and tissue-based bioassays are now routinely used for treatment planning.^3,4^ Here we present a strategy for extending this approach to the diagnosis and treatment of human spinal cord injury (SCI), a devastating and heretofore intractable condition characterized by injury heterogeneity and highly variable outcomes. Currently, SCI prognostics are based principally on acute evaluation of neurological status using sensory and motor exams including the American Spinal Injury Association (ASIA) grading system,^5^ which in the acute phase can be unstable and difficult to obtain, especially when patients are unresponsive or obtunded.^6-9^ Magnetic resonance imaging (MRI) provides invaluable information on severity and spinal cord level of injury, but is not always available and may be contraindicated for certain patients, e.g. those with penetrating metallic injuries.

The first attempts to discover SCI biomarkers of initial injury severity and long-term outcomes, date back four decades.^10-12^ Progress has been slow due at least in part to the diffuse regional presentation of acute SCIs and the difficulty of obtaining ultra-acute samples and patient data. Most attempts have used proteomics to identify serum and cerebrospinal fluid (CSF) biomarkers associated with injury severity. This approach depends upon measuring proteins associated with CNS damage (e.g. GFAP, neurofilament protein) released into the bloodstream, or on the peripheral cytokine response to CNS-injury-induced chemokines.^13-20^ Recent work shows promise, with several target molecules providing some useful predictive value,^21-26^ however, these circulating protein (and recently, RNA) markers are difficult to measure and subject to degradation. An alternative approach is to consider that circulating immune cells represent ‘sensors’ of CNS-injury-induced molecules, and that WBC transcriptomic changes provide a read-out of the complex peripheral immune response to the totality of signals associated with SCI over time. A recent study of WBCs in people with chronic SCI, for example, found reduced expression of genes associated with natural killer (NK) cell activity and increased expression of toll-like receptor and inflammatory cytokine genes^27^. These preliminary findings, although limited by a small number of cases, are consistent with observations that people with chronic SCI have suppressed immune responses and are more susceptible to infections.^28-30^

TRACK-SCI (“Transforming Research And Clinical Knowledge-SCI”) is a multicenter prospective clinical study focused on acute critical care variables (e.g. MRI, multiple physiological variables, time to surgery) and blood transcriptomics as indices of severity and predictors of outcome.^31-40^ SCI has a profound impact on circulating white blood cells (WBCs), inducing peripheral inflammation and WBC phenotypic changes in a dynamic cascade that is likely to reflect the biological features of the evolving CNS lesion.^41-44^ TRACK-SCI provides a large WBC transcriptomic dataset that will be useful for developing RNA-based blood biomarkers of injury and recovery that can be related to patient characteristics and multivariate outcomes. Utilizing advanced analytic methods, these blood biomarkers, along with other critical care variables, may be instrumental in the development of predictive algorithms for acute SCI treatment planning, to stratify patients for clinical trials and to predict long-term outcomes.

We are using deep RNA sequencing and advanced analytics to develop a blood RNA biomarker profile for acute SCI. Here, we report that early WBC transcriptomic signatures alone can accurately predict injury severity on the ASIA impairment scale (AIS). Further, these signatures provide novel biological data that should be useful in understanding mechanisms of injury and repair. Our findings provide proof of concept for the development of an accurate blood RNA biomarker of acute SCI severity. These data provide a strong rationale for expanding this work to include longitudinal multivariate analysis of gene expression patterns across injury severities, individual patient characteristics, and time, in order to provide a comprehensive description of evolving WBC gene expression patterns and their relationships to long-term outcomes.

## RESULTS

### TRACK-SCI patient accrual, data collection, and retention

TRACK-SCI protocols for patient enrollment and data collection have been described recently.^45^ Patients admitted to the emergency department at the Zuckerberg San Francisco General Hospital and Trauma Center (ZSFG) are recruited to the study and consented as soon as possible after admission. The TRACK-SCI protocol includes rapid pre-operative imaging, emergent transfer to the OR for decompression surgery as indicated, immediate blood collection and processing, followed by high-density ICU monitoring of vitals and daily sensorimotor and International Standards for the Neurological Classification of Spinal Cord Injury (ISNCSCI) exams (see **Fig S1**). To date, 179 participants with SCI have been enrolled. The current report is based on deep sequencing of RNA from acute blood samples from 38 subjects with SCI, 10 healthy uninjured controls (HC), and 10 trauma controls with non-CNS injuries (TC; see **Table 1**).

**Table 1.** Demographic data for patients in the analysis.

|  | HC<br>(n=10) | TC<br>(n=10) | SCI<br>(n=38) |
| --- | --- | --- | --- |
| **Sex** |  |  |  |
| Male | 8 (80.0%) | 6 (60.0%) | 25 (65.8%) |
| Female | 2 (20.0%) | 4 (40.0%) | 13 (34.2%) |
| **Race** |  |  |  |
| Asian | 4 (40.0%) | 2 (20.0%) | 10 (26.3%) |
| Black or African-American | 0 (0%) | 0 (0%) | 5 (13.2%) |
| Hispanic | 0 (0%) | 1 (10.0%) | 3 (7.9%) |
| White | 4 (40.0%) | 4 (40.0%) | 17 (44.7%) |
| Other | 0 (0%) | 3 (30.0%) | 0 (0%) |
| Unknown | 2 (20.0%) | 0 (0%) | 3 (7.9%) |
| **Age (years)** |  |  |  |
| Mean (SD) | 49.4 (12.2) | 41.7 (18.0) | 55.3 (20.0) |
| Median [Min, Max] | 51.0 [31.0, 67.0] | 42.0 [22.0, 78.0] | 53.5 [20.0, 89.0] |
| Missing | 1 (10.0%) | 0 (0%) | 0 (0%) |
| **Injury Severity Score (ISS)** |  |  |  |
| Mean (SD) | NA (NA) | 6.57 (4.35) | 27.8 (13.2) |
| Median [Min, Max] | NA [NA, NA] | 5.00 [1.00, 14.0] | 26.5 [9.00, 75.0] |
| Missing | 10 (100%) | 3 (30.0%) | 2 (5.3%) |
| **Prior CNS Pathology** |  |  |  |
| Yes | 0 (0%) | 0 (0%) | 12 (31.6%) |
| No | 10 (100%) | 10 (100%) | 21 (55.3%) |
| Unknown | 0 (0%) | 0 (0%) | 5 (13.2%) |
| **Concurrent TBI** |  |  |  |
| Yes | 0 (0%) | 0 (0%) | 8 (21.1%) |
| No | 10 (100%) | 10 (100%) | 28 (73.7%) |
| Unknown | 0 (0%) | 0 (0%) | 2 (5.3%) |
| **Time from blood draw (hours post injury)** |  |  |  |
| Mean (SD) | NA (NA) | 21.5 (5.75) | 30.3 (18.9) |
| Median [Min, Max] | NA [NA, NA] | 20.0 [15.0, 32.0] | 23.0 [5.00, 97.0] |
| Missing | 10 (100%) | 4 (40.0%) | 0 (0%) |

**Table 2.** Neurological examination of the SCI patients. To avoid the previously reported variability of the AIS grade during the first 2 days after SCI, we used the AIS grade assigned between days 3 and 10 post SCI in our analysis. 5 out of 38 SCI patients whose blood was sequenced did not have an examination during that time period and were therefore excluded from the analysis.

|  | A<br>(n=12) | B<br>(n=4) | C<br>(n=6) | D<br>(n=11) | Overall<br>(n=33) |
| --- | --- | --- | --- | --- | --- |
| **Level of injury** |  |  |  |  |  |
| Cervical | 2 (16.7%) | 3 (75.0%) | 3 (50.0%) | 10 (90.9%) | 18 (54.5%) |
| Thoracic | 8 (66.7%) | 1 (25.0%) | 1 (16.7%) | 0 (0%) | 10 (30.3%) |
| Lumbar | 1 (8.3%) | 0 (0%) | 0 (0%) | 1 (9.1%) | 2 (6.1%) |
| Unable to determine | 1 (8.3%) | 0 (0%) | 2 (33.3%) | 0 (0%) | 3 (9.1%) |
| **AIS actual or estimate** |  |  |  |  |  |
| actual | 7 (58.3%) | 1 (25.0%) | 3 (50.0%) | 9 (81.8%) | 20 (60.6%) |
| estimate | 5 (41.7%) | 3 (75.0%) | 3 (50.0%) | 2 (18.2%) | 13 (39.4%) |
| **Upper Extremities Motor Score**<br>**(out of 50 points)** |  |  |  |  |  |
| Mean (SD) | 39.9 (17.6) | 20.3 (25.7) | 26.0 (21.4) | 34.7 (14.4) | 34.0 (17.9) |
| Median [Min, Max] | 50.0 [0.00, 50.0] | 7.00 [4.00, 50.0] | 19.0 [9.00, 50.0] | 42.0 [12.0, 50.0] | 44.0 [0.00, 50.0] |
| Missing | 2 (16.7%) | 1 (25.0%) | 3 (50.0%) | 1 (9.1%) | 7 (21.2%) |
| **Lower Extremities Motor Score**<br>**(out of 50 points)** |  |  |  |  |  |
| Mean (SD) | 5.00 (15.8) | 5.00 (4.58) | 2.33 (3.21) | 46.2 (5.75) | 20.5 (23.1) |
| Median [Min, Max] | 0.00 [0.00, 50.0] | 6.00 [0.00, 9.00] | 1.00 [0.00, 6.00] | 49.5 [36.0, 50.0] | 6.00 [0.00, 50.0] |
| Missing | 2 (16.7%) | 1 (25.0%) | 3 (50.0%) | 1 (9.1%) | 7 (21.2%) |
| **Sensory Score (Touch)**<br>**(out of 112 points)** |  |  |  |  |  |
| Mean (SD) | 48.4 (19.2) | 98.0 (NA) | 24.5 (6.36) | 89.8 (23.7) | 63.1 (31.3) |
| Median [Min, Max] | 46.5 [15.0, 74.0] | 98.0 [98.0, 98.0] | 24.5 [20.0, 29.0] | 96.5 [60.0, 112] | 60.0 [15.0, 112] |
| Missing | 4 (33.3%) | 3 (75.0%) | 4 (66.7%) | 5 (45.5%) | 16 (48.5%) |
| **Sensory Score (Pain)**<br>**(out of 112 points)** |  |  |  |  |  |
| Mean (SD) | 49.0 (19.1) | 98.0 (NA) | 24.0 (12.7) | 81.5 (22.7) | 60.4 (28.6) |
| Median [Min, Max] | 47.0 [15.0, 74.0] | 98.0 [98.0, 98.0] | 24.0 [15.0, 33.0] | 74.5 [61.0, 110] | 61.0 [15.0, 110] |
| Missing | 4 (33.3%) | 3 (75.0%) | 4 (66.7%) | 5 (45.5%) | 16 (48.5%) |
| **Time of ISNCSCI (days)** |  |  |  |  |  |
| Mean (SD) | 5.08 (2.68) | 4.00 (2.00) | 3.83 (0.983) | 4.73 (1.79) | 4.61 (2.06) |
| Median [Min, Max] | 4.00 [3.00, 10.0] | 3.00 [3.00, 7.00] | 3.50 [3.00, 5.00] | 5.00 [3.00, 7.00] | 4.00 [3.00, 10.0] |

### The WBC transcriptome separates SCI patients from trauma and healthy controls

We isolated 4ml of peripheral blood from 38 enrolled patients within a few hours (**Fig S1, Table 1**) after SCI. The blood was immediately processed, and total RNA was isolated from WBCs. The same procedure was followed for 10 TCs and 10 HCs. After RNAseq, raw counts were produced and normalized, and a T-distributed Stochastic Neighbor Embedding (t-SNE) plot was created using the principal components responsible for 90% of the variance (**Fig 1A**). The three groups are clearly separated based on their transcriptomic status at the time of the blood draw. These results support our hypothesis that the transcriptome of the WBCs contains valuable information about the pathophysiological status of the patients and warrant a more sophisticated and deeper analysis to reveal more details. Next, we performed differential gene expression (DGE) analysis among the three groups, revealing 2,096 genes that were significantly altered (> 2-fold change, adjusted p-value < 0.05) only in the SCI population (**Fig 1B** and **S2**). We then queried, how many of those genes display an expression pattern that follows the injury severity levels of the AIS grade. Among the 2,096 genes that were differentially expressed after SCI, 197 of them showed directional expression patterns with SCI severity. 117 of them increased their expression with injury severity and 80 decreased their expression with injury severity (**Fig 1C**). Gene ontology enrichment analysis showed that processes involved in the immune response and cellular secretion and localization were the most highly enriched. (**Fig S3**).

**Figure 1.**
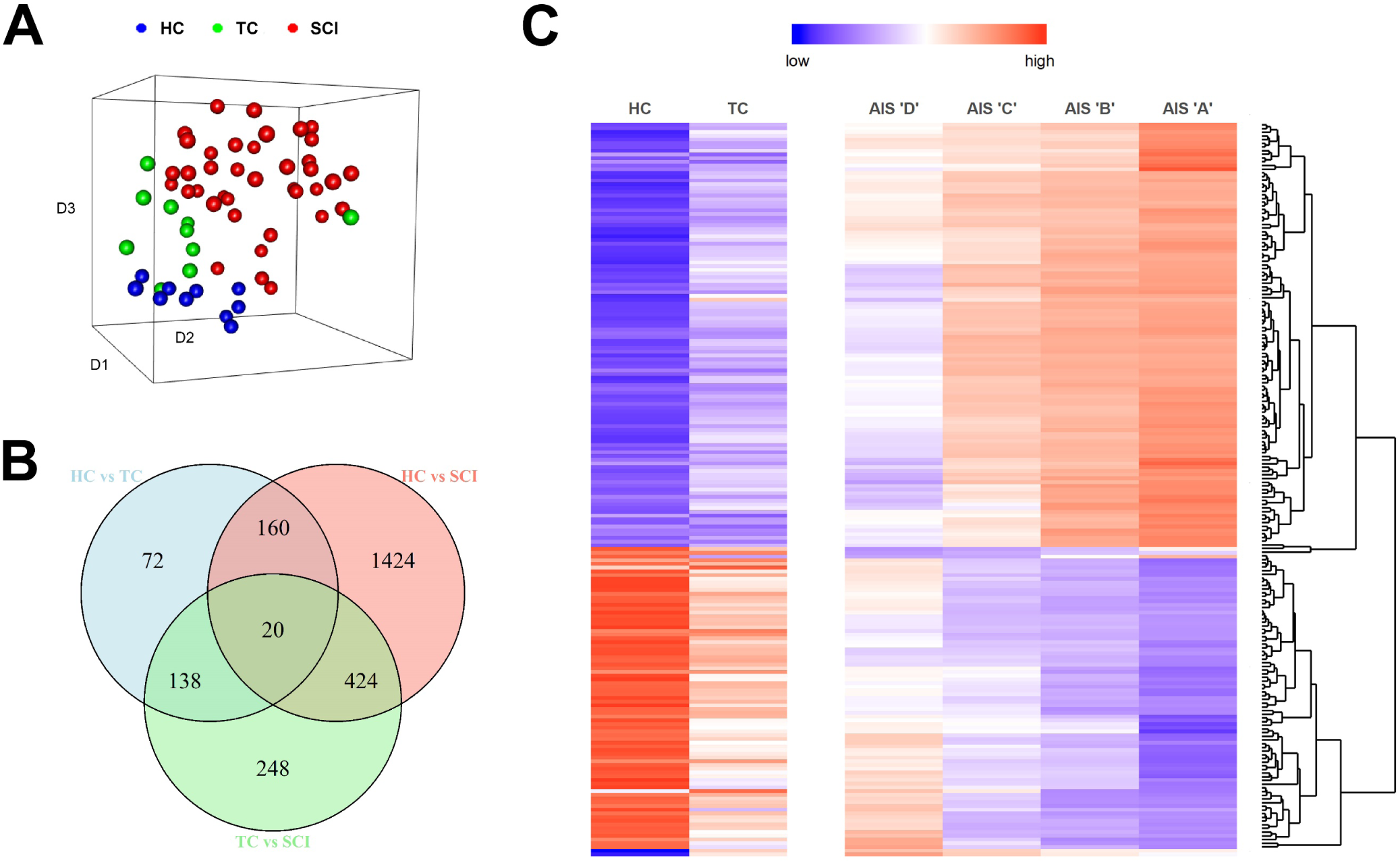
Spinal cord injury induces transcriptomic changes in white blood cells compared to healthy and non-CNS trauma controls. **A.** 3-dimension t-SNE plot. Each point on the plot represents one patient. The gene expression values of 17,500 transcripts were used in a principal component analysis, and the components that account for 90% of the variance were collapsed in the three dimensions of the t-SNE plot. The three groups (healthy controls [HC], trauma controls [TC], and spinal cord injury [SCI]) occupy different locations in the 3-dimensional space indicating that the transcriptomic signature alone is sufficient to separate them (HC = 10, TC = 10, SCI = 38). **B.** Differential gene expression analysis. The Venn diagram shows the intersection between differentially expressed genes for all three comparisons between HC, TC and SCI patients (fold-change > 2, adjusted p-value < 0.05). **C.** From the Venn diagram, we selected the genes that are only significantly changed after SCI and not in the event of trauma (1,424 + 424 + 248 = 2,096). Out of those 2,096 genes, 197 exhibit changes according to the AIS grade. The heatmap shows the expression pattern of these 197 genes. The upper part shows 117 genes whose expression increases as SCI severity increases and the bottom part shows 80 genes whose expression decreases as SCI severity increases (HC = 10, TC = 10, AIS D = 11, C = 6, B = 4, A = 12; AIS grade evaluated between 3 and 10 days post-SCI, and 5 patients did not receive an exam during that timeframe).

### Co-expression network analysis reveals gene modules that predict SCI severity

In order to study the modular organization of WBC transcriptomes in our sample cohort, we performed unsupervised gene co-expression network analysis^46,47^ and identified 16 modules (arbitrarily designated M1-M16) for which SCI patients’ combined expression was significantly different from both TCs and HCs (**Fig 2A**). These modules represent coherent transcriptomic signatures in WBCs that covary specifically as a result of SCI and are therefore targets for biomarker generation as well as potential indicators of underlying pathology and recovery. Among these, M13 exhibited the strongest correlation with AIS grade (Spearman rho = 0.82, p-value = 1.56 × 10^−14^). **Fig 2B-C** shows the details of top gene co-expression patterns for this module.

**Figure 2.**
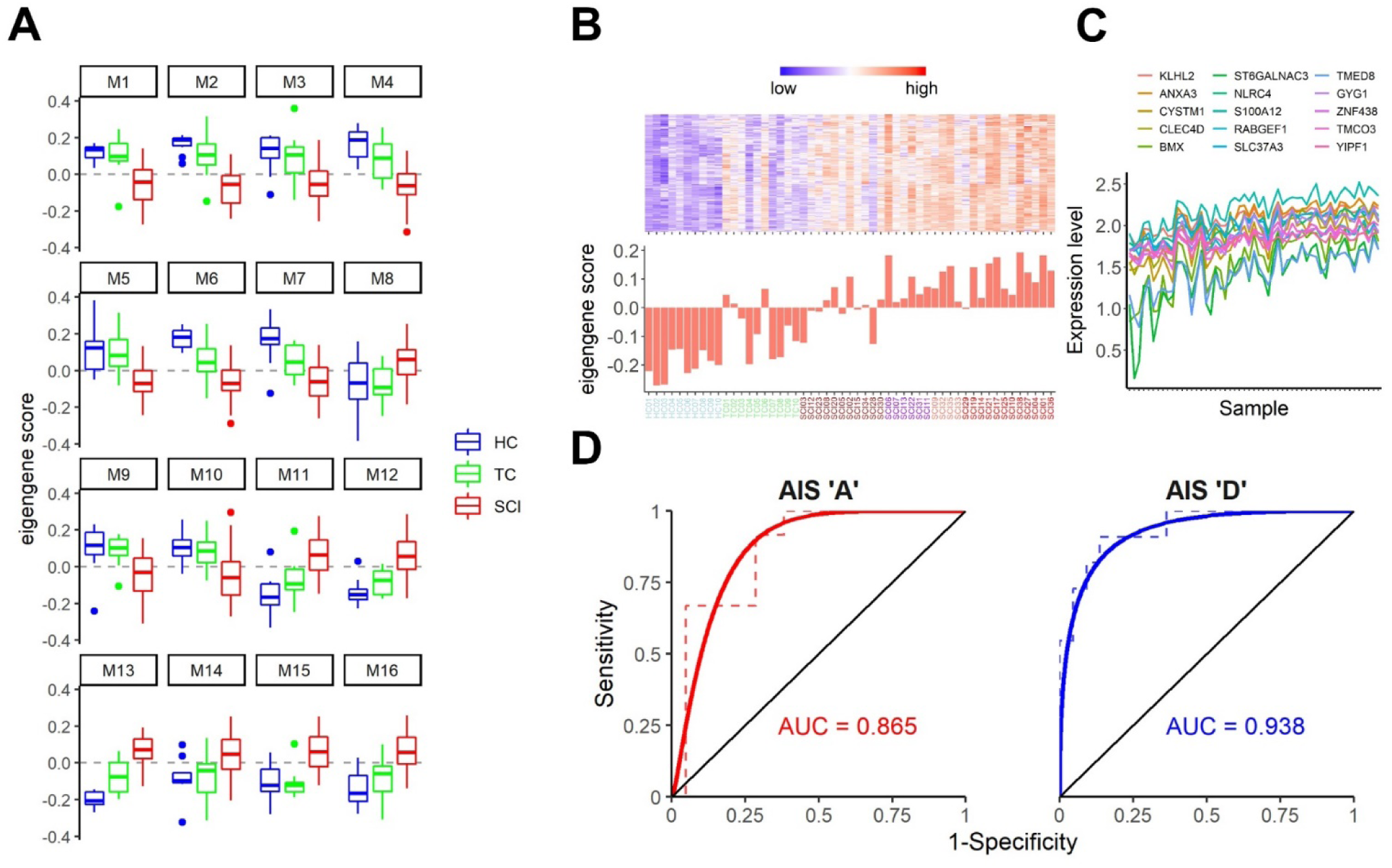
Gene co-expression network analysis reveals transcriptional modules in peripheral white blood cells that predict spinal cord injury severity. **A.** Analysis of module eigengene (PC1) scores by patient cohort reveals 16 SCI-specific gene co-expression modules following unsupervised gene co-expression network analysis. Some modules (e.g. M4) display a gradual change in gene expression, whereas in others (e.g. M1, M5) HCs and TCs are very similar to each other but different from SCIs. N = 10 for HCs and TCs and 38 for SCIs. **B – C.** The M12 module predicts AIS ‘A’ SCI patients with 83.3% accuracy. In B is the heatmap of the top-seeded genes for this module, and in C is the eigengene score for each one of the patients (and controls) for this module. The graph in C shows the expression levels of the top 15 genes of the M12 module across all 58 samples. As expected of the analysis, these top genes of the module exhibit a strong co-expression pattern. **D.** Receiver operating characteristic plots for the AIS ‘A’ against the remaining SCIs (left) and the AIS ‘D’ against the remaining SCIs (right). These plots show the strong predictive ability of our model for SCI patients with AIS ‘A’ and ‘D’. The AUC is 0.865 for the ‘A’ and 0.938 for the ‘D’. N = 12 ‘A’ vs 21 SCIs and 11 ‘D’ vs 22 SCIs (color scheme in x-axis labels in panel B is as follows: blue = HC, green =TC, brown = AIS ‘D’, purple = AIS ‘C’, salmon = AIS ‘B’, and red = AIS ‘A’).

Next, we wanted to determine whether any of these 16 modules, alone or in combination, could predict the initial SCI severity as indicated by the AIS grade assigned between days 3-10 after SCI. We therefore performed multinomial logistic regression with LASSO^48^ regularization to predict AIS grade using all 16 module eigengenes. We identified one gene module (M12; **Fig. S4**) that predicted AIS ‘A’ patients with 83.3% accuracy and a combination of five modules (M1, M5, M10, M13, and M16) that predicted AIS ‘D’ patients with 90.9% accuracy (**Fig. S4, Tables 3** & **S1**). Our cohort included too few patients with ‘B’ or ‘C’ classifications (n=4 and 6 respectively) to provide useful predictors of these grades (see supplemental files). Overall, our model shows 72.7% accuracy (p-value = 2.35 × 10^−5^). We proceeded to test the diagnostic value of the identified modules to detect AIS ‘A’ and ‘D’ patients in our cohort using a receiver operating characteristic (ROC) analysis. The area under the curve for AIS ‘A’ patients and AIS ‘D’ patients was 0.865 and 0.938, respectively, confirming that our model can predict these two injury severities with high sensitivity and specificity, despite small sample sizes (**Fig 2D**). Together these data show for the first time that WBC expression profiles can very accurately classify SCI patients based on AIS grade.

**Table 3.** Confusion matrix. This table known as a confusion matrix, shows the performance of our model. One can see that from the 12 AIS ‘A’ patients our model predicts 10 of them correctly (Sensitivity = 83.3%) but also misclassifies 5 more patients as ‘A’ (Specificity = 76.2%). For AIS ‘B’, ‘C’ and ‘D’ the sensitivity is 25%, 50% and 90.9% respectively, and the specificity is 100%, 100% and 81.2% respectively. Overall, the accuracy of our model is 72.7% (p = 2.35 × 10^−5^).

|  |  | Reference |  |  |  |
| --- | --- | --- | --- | --- | --- |
| Prediction |  | A | B | C | D |
|  | A | 10 | 2 | 2 | 1 |
|  | B | 0 | 1 | 0 | 0 |
|  | C | 0 | 0 | 3 | 0 |
|  | D | 2 | 1 | 1 | 10 |

### “Digital cytometry” and module enrichment analysis identifies WBC subtype differences between SCI and controls

To determine whether cellular composition varied among WBCs from our sample cohorts, we performed cell-type deconvolution with CIBERSORTx,^49,50^ which infers cell-type proportions in bulk tissues from global gene expression patterns. Applying this method to our samples revealed the relative proportions of 22 leukocyte subtypes. These “digital cell types” were then compared across groups (HC, TC and SCI) and AIS grade levels (**Fig. S5A-B**). Five cell types (neutrophils, resting NK cells, and resting CD4, naïve CD4 and gamma delta T cells) exhibited significantly different proportions among the 3 groups, but none of these were significant when AIS grades were compared.

To clarify which cell types contributed to the co-expression modules that predict SCI severity, we cross-referenced module composition with cell type-specific gene sets from published single-cell RNA-seq datasets.^49,51^ The M12 module, which predicted AIS ‘A’ injury severity, was significantly enriched with genes expressed by resting NK cells, mast cells, and CD66+ granulocytes (Fisher’s exact test p-values of 2.9 × 10^−7^, 5.19 × 10^−7^, and 7.81 × 10^−6^ respectively). The M13 module, which predicted AIS ‘D’ severity, was also significantly enriched with CD66+ granulocytes (p = 1.45 × 10^−8^) (see supplemental files). These data suggest that expression changes in WBCs associated with SCI can be further subdivided into specific contributions from distinct cell types, which could lead to refined assays and predictors.

## DISCUSSION

Early biomarkers of SCI could enable more efficient and personalized clinical treatments, as well as better stratification of patients for clinical trials, since early clinical evaluations often lack reliability.^7,19^ In contrast to existing biomarkers that require rapid access to complex machinery (e.g. MR scanning), fluid biomarkers are more accessible and can provide novel insights about systemic responses to SCI. Although previous studies have proposed large structural proteins in blood serum/plasma and CSF as fluid biomarkers of SCI,^14,23,52^ these proteins are susceptible to proteases and degrade rapidly, rendering measurements of concentration variable and time sensitive. More recent studies of circulating miRNAs as biomarkers are promising since miRNAs are not as sensitive to degradation due to their small size (22nt) and the fact that most are protected inside exosomes.^24,25,53,54^

Our approach differs substantially from previous efforts to identify fluid biomarkers of SCI. First, we have analyzed potential biomarkers that are ‘safely housed’ in their cell of origin at the time of collection, as opposed to free-floating molecules in CSF or blood. Second, instead of pre-selecting candidate biomarkers in advance, we have performed a high-throughput analysis of 17,500 transcripts from WBCs isolated from each patient. This high-dimensional readout of the immune response during the acute phase of the injury provides important information about how the periphery affects the progress of the central lesion and may lead to new hypotheses and targets for intervention. Our results indicate that global gene expression patterns in WBCs can reliably distinguish SCI patients from healthy and non-CNS trauma controls. Moreover, when overlaying these gene expression patterns with the widely used AIS injury grade classification system, we identified 197 genes whose expression levels changed with increasing injury severity and may serve as novel fluid biomarkers of SCI. The list of these 197 genes includes a number of genes that seem appropriate, as well as some whose functions are yet to be delineated, which might provide clues to new therapeutic targets (see supplemental files). However, there is growing evidence (mainly from cancer research) to suggest that single genes do not usually make good biomarkers.^55^ Of course, genes do not work in isolation, and there is overwhelming evidence for reproducible transcriptional covariation in blood and other tissues, suggesting a modular organization to genomic function. These modules are not always neatly captured by existing gene ontologies, hence the need to perform unsupervised co-expression analysis. Multivariate analysis revealed 16 gene co-expression modules associated with SCI, a subset of which were able to predict AIS ‘A’ and AIS ‘D’ SCI patients with impressive accuracy. From a clinical perspective this finding can be a ‘game-changer’ considering that in many instances, several hours and logistically challenging infrastructure (e.g. MRI) is needed even for confirmation of the SCI, let alone for determining the degree of severity.

Another goal of our study is to predict long-term outcomes (e.g. the AIS grade at 6- and 12-months post SCI). We are collecting longitudinal data from enrolled patients and in the future will be able to extend our analysis in this fashion. Moreover, due to the well-known limitations of the AIS grade scale, additional outcome measures (e.g. upper and lower extremity motor scores and sensory scores from the ISNCSCI neurological examination) could offer more detailed insights into patients’ conditions and enable more accurate predictions. As we increase the number of enrolled patients and sequenced blood samples, such analyses will be feasible.

Although we have demonstrated proof-of-concept for our methodology in a sample of 38 SCIs, TRACK-SCI continues to enroll patients, with 179 participants to date. By combining multivariate and longitudinal analysis of WBC transcriptomes with detailed clinical information, we seek to create a novel framework for diagnosing SCI severity and predicting outcomes based on the systemic immune response. Such a framework could eventually replace the ASIA grading system or be used in combination to allow more precise clinical decisions. Additional studies with larger sample sizes will be required to validate these findings and move towards a practical RNA-based blood biomarker. Furthermore, it should be possible to determine whether diagnostic patterns reflect WBC responses to specific CNS-injury-induced signals such as CNS protein products and/or signaling molecules such as chemokines. The clear differentiation between expression profiles from trauma controls and SCI suggests that WBC transcriptomes contain latent information specific to CNS injury, raising the possibility that WBC RNA profiles may also respond to treatments that mitigate secondary damage or are associated with recovery of function. If so, the routine analysis of RNA expression profiles in blood may provide both a practical clinical tool and a new window into the biology of human CNS injury and its treatment.

## Acknowledgements

The current work was supported by grants from the Department of Defense (W81XWH-13-1-0297 and W81XWH-16-1-0497) and Craig H. Neilsen Foundation (UCSF SCI Center of Excellence special project award) to M.S.B. and an individual grant from Wings for Life (WFL-US-07/18) to N.K. The authors would also like to thank the Center for Advanced Technology (CAT) at UCSF for providing consultation and critical equipment for the completion of this study.

## Author contributions

Conception and design: N.K., A.R.F., M.C.O., J.C.B., M.S.B.; Data collection: N.K., X.D.F., L.H.T., R.E.T., D.D.H., L.U.P., V.S., W.D.W., S.S.D.; Data curation and analysis: N.K., A.T.E., P.G.S., J.R.H., A.C., X.D.F., L.H.T., R.E.T., A.R.F., M.C.O., J.C.B., M.S.B.; Manuscript preparation: N.K., A.T.E., P.G.S., M.C.O., J.C.B., M.S.B.; Manuscript revision: all authors; Funding acquired: N.K., M.S.B.

## Competing interests

The authors declare no competing interests.

## Data availability statement

All data generated or analyzed during this study are included in this published article and its supplementary information files.

## MATERIALS AND METHODS

### TRACK-SCI patients and controls enrollment

All procedures for this study were conducted with the approval of the Human Subjects Review Boards at the University of California at San Francisco (UCSF) and the U.S. Department of Defense Human Research Protection Office. All English and non-English speaking patients who presented to the emergency department (ED) and were diagnosed with a traumatic SCI were initially eligible for the study. Patients who were < 18 years old, in-custody, prisoners, pregnant, or on medically indicated psychiatric hold were excluded. Informed consent was sought for all patients. For patients who were unable to sign for themselves due to their injury, a witness unaffiliated with the study was present throughout the consenting process and signed on the patient’s behalf. Patients incapable of consenting themselves were initially enrolled via a legally authorized representative (LAR; next of kin) or another suitable surrogate when one was available, then later approached for patient consent. Patients and surrogates had the option to participate in all or some of the following study portions: blood draws, ISNCSCI exams, and/or follow-up assessments. Patients were compensated ($50) after each time point (hospital stay, 3-month phone call, 6-month in-person visit, 12-month in-person visit) for a total of $200.

Non-SCI subjects were either healthy controls (n=10) or trauma controls (n=10). Healthy controls were recruited using IRB-approved recruitment flyers posted at ZSFG and from friends and family of enrolled SCI patients. Subjects contacted study coordinators, were interviewed and consented, and provided basic demographic information and biospecimens. Trauma controls were recruited from ED patients with traumatic but non-CNS injuries. The same basic demographic and biospecimen data were collected for these patients as SCI patients for comparison purposes patients except for multiple blood samples per patient. No monetary compensation for participation was provided for the control subjects.

### Patient data collection

The foundation of the TRACK-SCI database is the NINDS-recommended common data elements (CDEs).^56^ Core CDEs are data elements that all SCI studies are strongly encouraged to use in collection of basic participant information. Additional measures from the International Spinal Cord Society (ISCoS) were also used. Data collection domains include demographic, clinical, radiologic, and functional outcome measures. All data collected from these CDEs were housed in a Research Electronic Data Capture (REDCap)^57,58^ database and include more than 21,000 data fields including additional institutional variables, calculated fields, repeated measures, date/time stamping of measures, and completion status log. Upon admission to the inpatient service, another 19,148 data fields regarding trauma characteristics, injury severity, blood pressure management, operating room procedures, interventions, hospital outcomes, high-frequency operating room vital signs, as well as motor-sensory exams and pain questionnaires are obtained from both paper and electronic medical records as well as participant interview. REDCap is in full compliance with Health Insurance Portability and Accountability Act (HIPAA) security standards for protection of personal health information (PHI). The following CDE categories comprised the demographic and clinical data domain: (1) demographics, (2) health history, (3) injury-related events, (4) assessments and examinations. A total of 229 variables concerning patient demographics, medical history, and consent/contact information were collected through abstraction from electronic medical record systems and participant interviews.

The International Standard for Neurological Classification of Spinal Cord Injury (ISNCSCI)^59^ was used to assess motor and sensory function, and group patients by injury severity based on the ASIA impairment scale (AIS) which ranges from A (most severe – complete) to E (not impaired). ISNCSCI exams were conducted by trained personnel who completed the ASIA International Standards Training E Program (InSTEP) and in-person training. ISNCSCI exams were performed on all patients during the initial admission, either as part of clinical care if the treating provider completed InSTEP training, or separately for the research study if the ISNCSCI was not performed for clinical purposes. Occasionally, an ISNCSCI was not performed or not completed during the admission, usually because the patient was excessively sedated and could not participate in the exam. In the case of incomplete ISNCSCI exams, the assessor gave an estimated AIS grade based on the collected data and the overall clinical picture of the patient.

If possible, patients completed examinations at regular intervals including admission (day 0 = 0-23 hours from injury), every 24 hours until post-injury day 7, discharge, 6-month follow-up (+/− 2 weeks), and 12-month follow-up (+/− 2 weeks). All ISNCSCI exams results were included in the REDCap database.

### Biospecimen collection

Two blood samples were collected, one, 4ml, for total RNA extraction from white blood cells (WBCs) and the other, 6ml, for serum isolation. Samples were aliquoted and frozen at −80°C within 1 hour of collection. To preserve WBC concentrations, 7.2mg K2-EDTA vacutainer tubes were used for WBCs collection and subsequent RNA extraction, instead of K3-EDTA vacutainer tubes. The second blood sample was collected in 6 ml Z serum clot activator vacuette tubes for serum extraction. To prevent reduction in sample volume, externally threaded cryovials were used for serum storage. Serum was divided into multiple 500 *µ*l aliquots for storage. An inventory system was developed to track time of collection, processing and storage for all biospecimens.

### WBC isolation and RNA extraction

Blood was centrifuged at 1,500 rpm for 15 minutes at room temperature within 0-15 minutes from blood draw. Then, the interface layer (buffy coat) was carefully aspirated with a pipette and placed in 10mL of 1X solution of Red Blood Cell Lysis Buffer (BioLegend) for 15 minutes in the dark. After the 15-minute incubation, the solution was centrifuged at 1,500 rpm for 10 minutes. The supernatant was discarded, and the cell pellet resuspended in 1mL of TRIZOL (Ambion) and either stored at −80°C or immediately processed for RNA extraction. Total RNA from WBCs was extracted using the TRIZOL method. The RNA yield was between 15 and 25μg per 5ml peripheral blood. 1μg of the total RNA was then used for generating the Illumina cDNA library, which was used for the downstream RNAseq.

### RNA sequencing

1μg of total RNA was used for the library synthesis. cDNA libraries were synthesized using Illumina’s TruSeq Stranded Total RNA with Ribo-Zero Globin kit. The kit depletes ribosomal RNA which makes up more than 90% of total RNA, and globin mRNA that is present in very high levels in blood total RNA. The libraries were quantified using a Thermo Scientific Nanodrop 2000c spectrophotometer, and their quality and average fragment size was assessed using Agilent’s DNA 1000 kit and Agilent’s 2100 Bioanalyzer. After quantification, equal amounts of 10 libraries, each one with different barcoded adapters, were pooled together to be sequenced in one lane of the Illumina’s HiSeq4000 sequencer. Based on the specifications of the HiSeq4000 and our sample pooling per lane, we aimed to get about 40 million reads per sample which has been shown to be sufficient to reveal the vast majority of the differentially expressed genes of a well-annotated genome.^60^ The sequencing output of our samples can be seen in **Table S2**.

### Bioinformatic analysis

Data analysis was performed in R^61,62^ using the statistical packages that are specifically mentioned below as well as the packages dplyr^63^, ggplot2^64^, cowplot^65^, table1^66^, rgl^67^, PCAtools^68^, magick^69^, EnhancedVolcano^70^, VennDetail^71^, Rtsne^72^, and ggdendro.^73^ The raw reads of the fastq files were tested for quality control using the FastQC software^74^ and were then aligned to the human reference genome (hg38 from UCSC) using the software TopHat2.^75^ After the alignment, we used the featureCounts^76^ program to summarize the gene counts, and then the programs edgeR^77^ and limma^78^ for differential gene expression analysis through linear modeling. Gene ontologies enrichment analysis was performed with the GOrilla^79^ tool and the visualization of the enriched GO terms with the tool ReviGO.^80^

### SCI-specific differentially expressed genes and neurological outcome

Preliminary examination of the data using network methods^81^ revealed several samples as outliers as well as library and sequencing batch effects. Using the ComBat^82^ function of the sva^83^ package in R to target the library batch effect we were able to remove both batch effects and as a result no sample appeared as an outlier anymore. We performed differential gene expression analysis among the three groups (HC, TC, and SCI) and after the intersection of the three comparisons (HC vs TC, HC vs SCI, and TC vs SCI; fold change > 2 and adjusted p-value < 0.05) we selected only the genes that significantly and specifically changed their expression after SCI (n = 2,096). In order to examine the relationship between the gene expression changes and injury severity, we grouped the SCI patients based on the assigned AIS grade given after a neurological examination between days 3 and 10 at the hospital. From the 38 SCI patients whose transcriptome was sequenced, 5 did not have an AIS grade during that time window and hence were removed from this part of the analysis. We then averaged the expression levels of each gene per group and queried for genes which exhibit a stepwise increased or decreased expression as the SCI severity increased. That query resulted in 197 genes shown in Fig 1C. The complete gene list can also be found in the supplemental files.

### Gene co-expression network analysis and identification of SCI-specific gene modules

The normalized expression matrix that was generated for the differential gene expression analysis with edgeR was used as a template for the unsupervised gene co-expression network analysis.^46,47^ We built a series of gene co-expression networks and identified one that included the gene module with the highest Spearman correlation to AIS grade. That network contained 57 gene modules. Using one-way ANOVA with Tukey’s multiple comparison correction we identified 17 gene modules that were highly specific for SCI (significantly different from both HC and TC; adjusted p-value < 0.05). One of these modules contained genes annotated to be involved in rRNA processes, which likely represented an artifact, and was eliminated from the subsequent analysis. The 16 remaining SCI-specific modules (M1 – M16; **Fig 2A**) were used to create a predictive model of SCI severity using multinomial logistic regression.

### Multinomial logistic regression with regularization

We generated a predictive model of SCI severity using the eigengenes (first principal components)^84^ of the 16 SCI-specific gene modules as predictors. AIS at discharge from the hospital was used as the target outcome variable in a multinomial logistic regression model. In order to deal with the high number of predictors (16 modules), LASSO regularization was applied, using leave-one-out cross-validation to determine the regularization parameter (λ). The final model was chosen with λ producing a model with a misclassification error at one standard deviation from the minimal misclassification error. The model was specified using the *glmnet*^*48*^ and the *glmnetUtils*^*85*^ packages in R. The model was assessed by confusion matrix metrics (**Tables 3** & **S1**) of internal prediction obtained using the *caret* R package.^86^ Overall accuracy (percentage of correct classification) was 72.7% with a 95%CI of 54.5-86.7%, resulting in significant accuracy (p-value<0.0001) against random classification (No Information Rate of 36.4%). The uniform weighted overall accuracy (accounting for class unbalance) was 62.3%, with accuracy for AIS ‘A’ = 83.3%, AIS ‘B’ = 25%, AIS ‘C’ = 50%, and AIS ‘D’ = 90.9%. Receiver operating characteristic (ROC) curves for each AIS class were obtained by binarizing the problem (e.g. for AIS ‘A’, A = 1; B, C, D = 0) and re-running the model as a binary classification. The curves were obtained using the *roc()* function of the *pROC* R package^87^ and smoothing transformation was applied to each ROC curve using the *smooth()* function of the *pROC* R package.

### CIBERSORTx and module enrichment analysis

CIBERSORTx^50^ is a machine learning algorithm that uses deconvolution methods to infer the proportions of cell types from gene expression patterns in bulk tissues. It is called digital cytometry because it performs an analogous function to regular flow cytometry without the need to physically isolate cells. We used the CIBERSORTx tool (https://cibersortx.stanford.edu/index.php) on all 58 of our samples and cross-referenced it with the LM22 signature^49^ using 100 permutations. LM22 is a validated leukocyte gene signature matrix containing 547 genes that distinguish 22 human hematopoietic cell types. The output of the algorithm is the relative abundance of each of the 22 subtypes for all samples (**Fig S5A**). For enrichment analysis, modules were defined as all unique genes with positive *k*_ME_ values (Pearson correlation coefficients of module eigengenes)^84^ that were significant after applying a Bonferroni correction for multiple comparisons (p < 0.05 / (# genes × # modules)). If a gene was significantly correlated with more than one module eigengene, it was assigned to the module for which it had the highest *k*_ME_ value. Enrichment analysis was performed for each gene set of interest with published human RNA-seq datasets,^49,51^ using a one-sided Fisher’s exact test as implemented by the fisher.test R function.

---

**Supplementary Figure 1.**
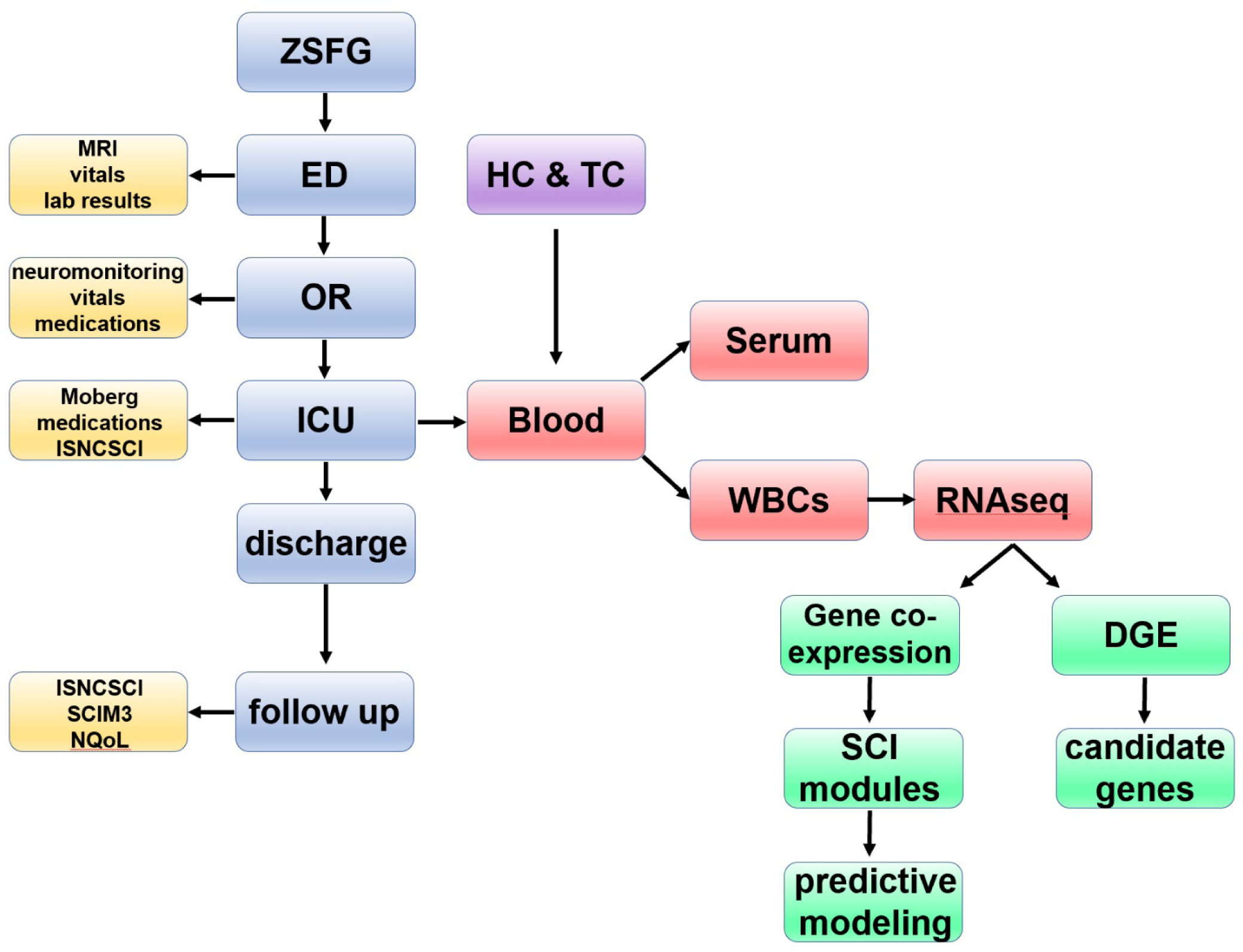
Flowchart of patient enrollment, data acquisition and analytic pipeline. In TRACK – SCI, as soon as a confirmed SCI patient is admitted and consents to participate in the study, our team collects clinical data during all stages of the hospital stay and at 3, 6 and 12 months post-injury (in total more than 22,000 data points for each patient). Blood is also collected as early as possible after hospital admission (day 0) and at days 1, 2, 3 and 5 as well as at 6- and 12-months post injury. After the blood draw, white blood cells are isolated, and RNA is extracted for RNAseq. The RNAseq data from SCI patients along with RNAseq data from Healthy and Trauma Controls are analyzed using both supervised and unsupervised methods with the goal of creating a predictive model for injury severity.

**Supplementary Figure 2.**
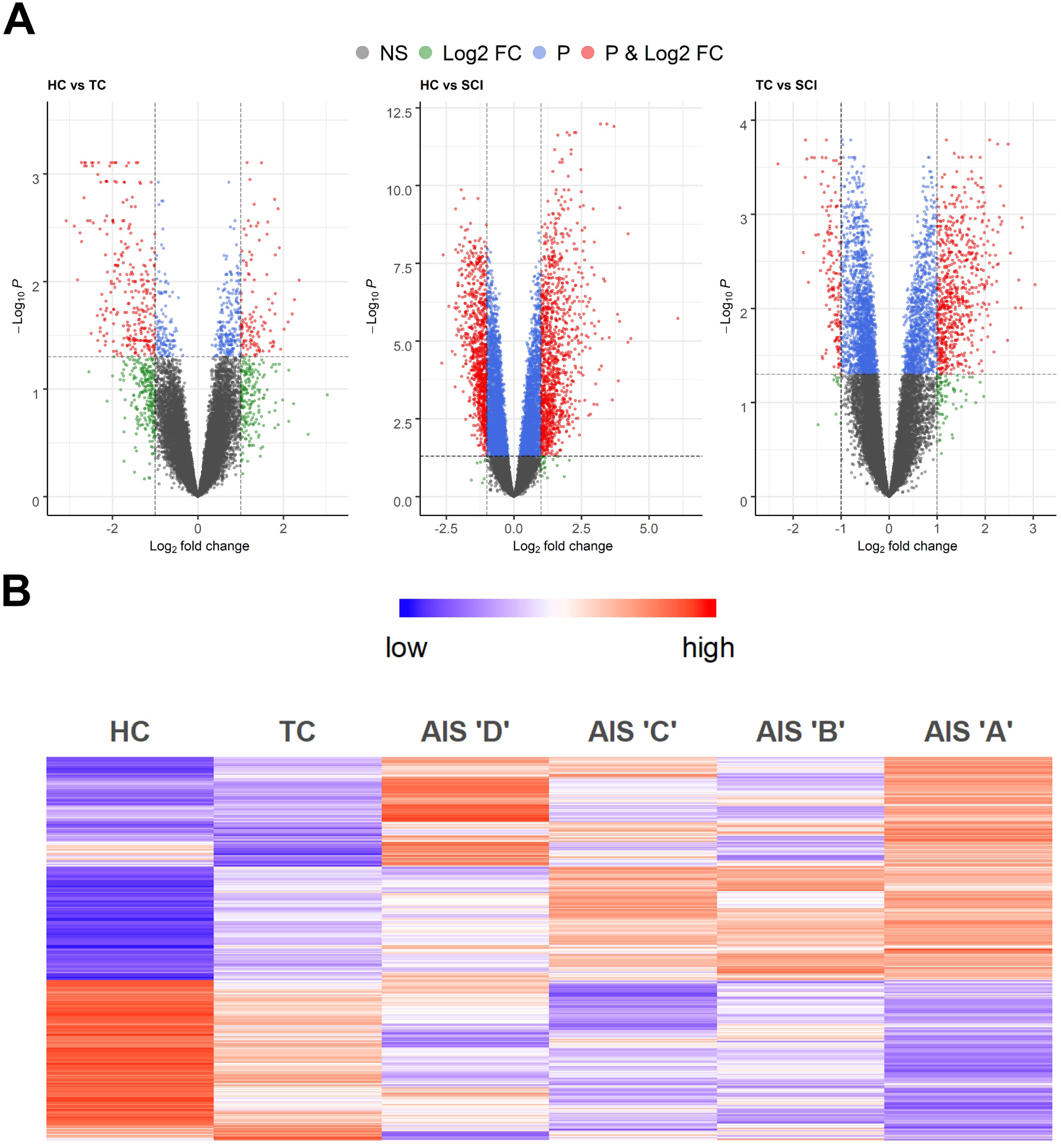
Differential gene expression analysis of SCI patients vs. healthy and trauma controls reveals many genes induced specifically upon SCI. **A.** Volcano plots of the 3 comparisons between Healthy Controls (HC), Trauma Controls (TC) and SCI patients (SCI). **B.** Heatmap of the 2,096 differentially expressed genes after SCI but not trauma (fold-change > 2, adjusted p-value < 0.05; HC = 10, TC = 10, AIS D = 11, C = 6, B = 4, A = 12).

**Supplementary Figure 3.**
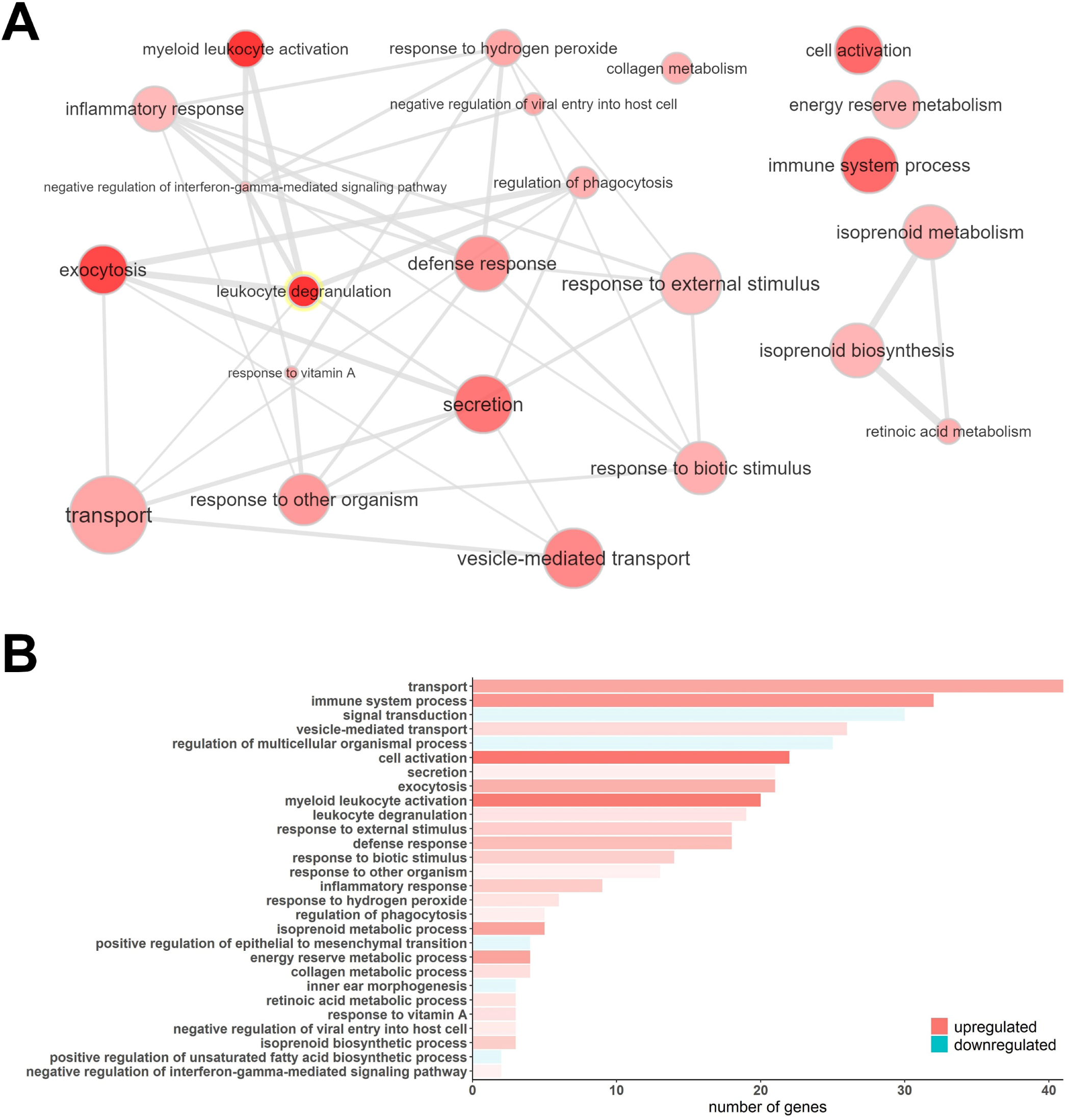
Gene Ontology (GO) enrichment analysis of the directional with SCI severity differentially expressed genes, suggests an important role of inflammation and cellular transport and localization in classifying SCI patients. **A.** Visualization of the enriched GOs of the genes that increase their expression as the AIS grade increases. The bubble color shading indicates the p-values (stronger shading = lower p-value) and the bubble size the frequency of the GO in the underlying GO Annotation database. The lines link highly similar GO terms and the width of the line indicates the degree of similarity. **B.** The bar plot shows the number of differentially expressed genes in each one of the significant GO terms. The shade of each bar indicates the p-value (stronger shading = lower p-value).

**Supplementary Figure 4.**
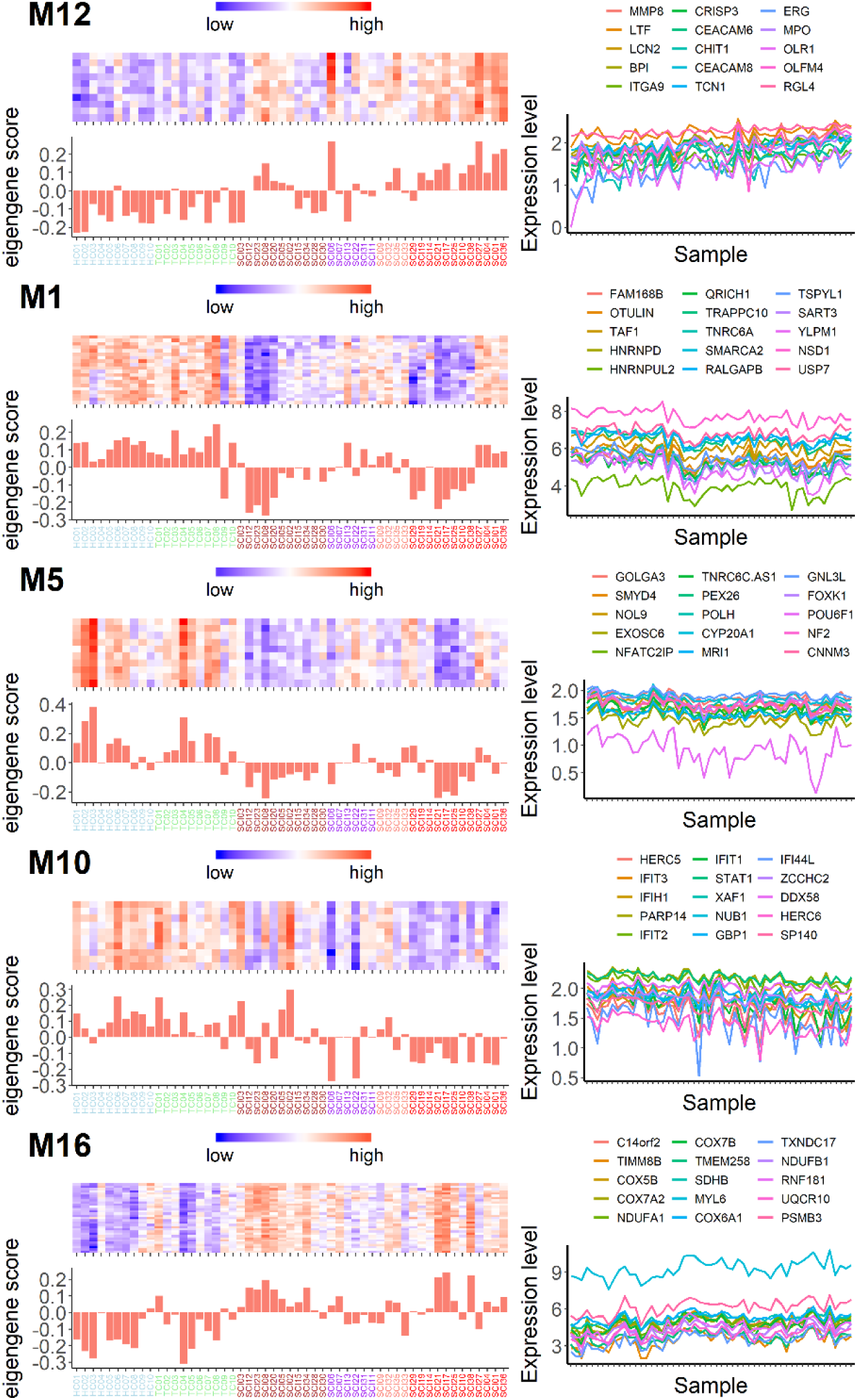
Multinomial logistic regression identifies specific gene modules with the capacity to accurately predict AIS ‘A’ and AIS ‘D’ SCI patients. For AIS ‘A’ SCI patients one gene module (M12) is sufficient to predict the injury class with 83.3% accuracy. Interestingly, five gene modules (M13 in Fig.2B-C, M1, M5, M10 and M16) are required to predict AIS ‘D’ SCI patients with an impressive 90.9% accuracy. For each one of the modules in this figure, on the left is a heatmap with the top-seeded genes for the module and the eigengene score for each patient (and control); on the right are the expression patterns (in arbitrary units) of the top 15 genes with the highest correlation to each module eigengene (color scheme in x-axis labels is as follows: blue = HC, green =TC, brown = AIS ‘D’, purple = AIS ‘C’, salmon = AIS ‘B’, and red = AIS ‘A’).

**Supplementary Figure 5.**
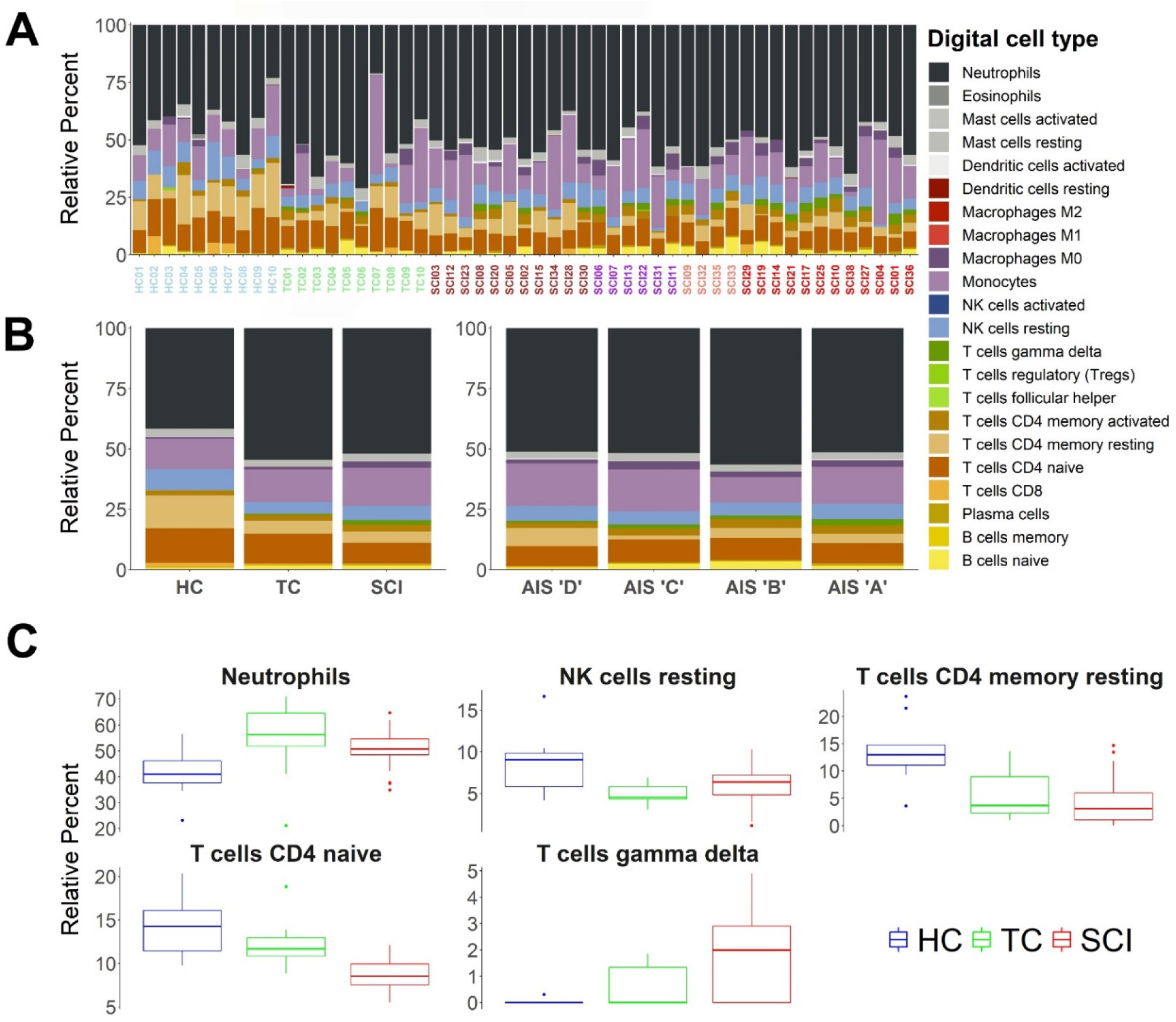
Digital cytometry using CIBERSORTx measures relative abundance of 22 distinct leukocyte subtypes in SCI and control patients. We used a recently created machine learning algorithm (CIBERSORTx) which uses deconvolution methods to infer cell type proportions based only on gene expression patterns. We cross-referenced the transcriptomes of all SCI patients and controls with the leukocyte gene signature matrix (LM22) and estimated relative abundances for 22 leukocyte subtypes. **A.** The stacked bar plots show the relative abundance of the 22 digital cell types for each of the SCI patients and controls. **B.** Left shows group averages and right shows AIS grade averages. **C.** One-way ANOVA for each “digital” cell type showed statistically significant differences for neutrophils, resting NK cells, CD4 resting T cells, CD4 naïve T cells, and gamma delta T cells with adjusted p-values < 0.05. Tukey’s test showed that for CD4 naïve and gamma delta T cells the SCI group is significantly different from both control groups. No statistically significant difference was identified among AIS grades (color scheme in x-axis labels in panel A is as follows: blue = HC, green =TC, brown = AIS ‘D’, purple = AIS ‘C’, salmon = AIS ‘B’, and red = AIS ‘A’).

**Table S1.**
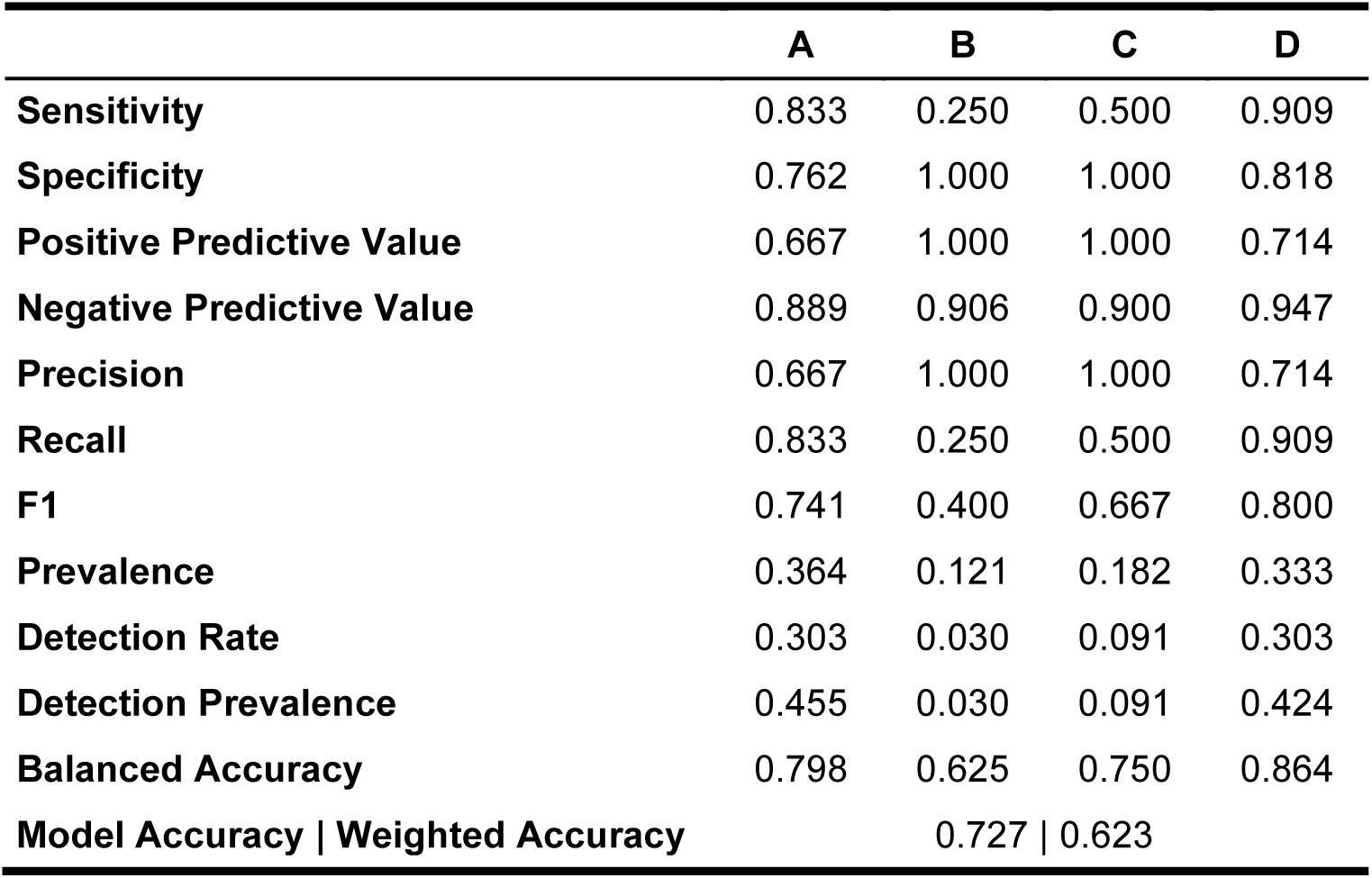
Predictive model summary statistics.

**Table S2.** Sequenced samples output.

|  | HC<br>(N=10) | TC<br>(N=10) | SCI<br>(N=38) | Overall<br>(N=58) |
| --- | --- | --- | --- | --- |
| **Number of reads (in millions)** |  |  |  |  |
| Mean (SD) | 45.7 (5.52) | 38.6 (5.10) | 41.0 (7.57) | 41.4 (7.13) |
| Median [Min, Max] | 45.9 [38.5, 56.6] | 38.9 [27.6, 44.5] | 40.5 [26.8, 73.4] | 40.9 [26.8, 73.4] |
| **% Perfect barcode** |  |  |  |  |
| Mean (SD) | 97.5 (0.593) | 96.7 (1.09) | 97.5 (0.696) | 97.4 (0.815) |
| Median [Min, Max] | 97.3 [96.8, 98.4] | 96.6 [95.2, 98.4] | 97.4 [96.1, 98.7] | 97.3 [95.2, 98.7] |
| **% One mismatch barcode** |  |  |  |  |
| Mean (SD) | 2.48 (0.593) | 3.33 (1.09) | 2.47 (0.696) | 2.62 (0.815) |
| Median [Min, Max] | 2.72 [1.60, 3.25] | 3.42 [1.59, 4.84] | 2.56 [1.28, 3.87] | 2.71 [1.28, 4.84] |
| **Yield (Mbases)** |  |  |  |  |
| Mean (SD) | 2320 (288) | 1940 (253) | 2070 (387) | 2090 (366) |
| Median [Min, Max] | 2320 [1950, 2890] | 1960 [1410, 2270] | 2030 [1340, 3740] | 2050 [1340, 3740] |
| **% >= Q30 bases** |  |  |  |  |
| Mean (SD) | 97.5 (0.452) | 97.8 (0.226) | 97.7 (0.358) | 97.7 (0.362) |
| Median [Min, Max] | 97.6 [96.9, 98.0] | 97.9 [97.5, 98.0] | 98.0 [96.8, 98.1] | 97.9 [96.8, 98.1] |
| **Mean Quality Score (max = 40)** |  |  |  |  |
| Mean (SD) | 39.5 (0.133) | 39.6 (0.0529) | 39.6 (0.0992) | 39.6 (0.101) |
| Median [Min, Max] | 39.6 [39.4, 39.7] | 39.6 [39.5, 39.7] | 39.7 [39.3, 39.7] | 39.6 [39.3, 39.7] |

## Notes

### Competing Interest Statement

The authors have declared no competing interest.

